# An *in-vitro* BBB-on-a-chip open model of human blood-brain barrier enabling advanced optical imaging

**DOI:** 10.1101/2020.06.30.175380

**Authors:** Mootaz Salman, Graham Marsh, Ilja Küsters, Matthieu Delincé, Giuseppe Di Caprio, Srigokul Upadhyayula, Giovanni de Nola, Ronan Hunt, Kazuka G. Ohashi, Fumitaka Shimizu, Yasuteru Sano, Takashi Kanda, Birgit Obermeier, Tom Kirchhausen

## Abstract

We describe here the design and implementation of an *in-vitro* BBB-on-a-chip open model system capable of reconstituting the microenvironment of the blood brain barrier. This system allows controlled unidirectional flow of nutrients and biologicals on the lumen of the artificial microvessel. This BBB-on-a-chip is suitable for high resolution electron microscopy and it is amenable for quantitative 3D live fluorescence imaging using spinning confocal disk or lattice light sheet microscopy (LLSM) to follow, for example the transcytosis across the BBB-like barrier of fluorescently-tagged biological, viruses or nanoparticles.

## INTRODUCTION

The blood–brain barrier (BBB) is a unique and highly selective vascular interface that separates the peripheral blood circulation from the neural tissue in order to maintain a homeostatic microenvironment within the central nervous system (CNS) that allows the neuronal network to function properly (Abbott et al., 2010; Park et al., 2019).

The BBB is a complex vascular structure with specialized endothelial cells as its core element, surrounded by extracellular matrix (ECM) and supporting cells, such as astrocytes and pericytes (Greene and Campbell, 2016). Brain microvascular endothelial cells (BMVECs) that line the capillaries of the BBB are of crucial physiological importance since they tightly control the molecular and cellular flux between the blood and the brain, thereby regulating regional changes in nutrients and oxygen levels (Daneman, 2012), maintaining brain energy levels (Bordone et al., 2019) and mediating the local immune response in the CNS (Abbott et al., 2010). BMVECs differ from those found in peripheral vasculature as they have no fenestration and exhibit restricted paracellular passage for water and hydrophilic solutes due to the presence of a unique array of tight junctions and adherens junctions between adjacent endothelial cells (Greene and Campbell, 2016). Moreover, BMVECs have specialized transcellular transport mechanisms ensuring only wanted substances being actively delivered to the brain, and have shown to express a number of broad-spectrum efflux pumps on their luminal surface which severely limit the uptake of lipophilic molecules, including small molecule drugs, from the blood through the endothelium into the CNS. These characteristic anatomical and functional features of the BBB determine its crucial protective role for the CNS (Mahringer et al., 2011; Shawahna et al., 2011).

However, these highly selective barrier properties also extremely limit the therapeutic efficacy of drugs and hinder the treatment of neurological diseases such as Alzheimer’s disease, multiple sclerosis, Parkinson’s disease, HIV infection and brain tumors (Pardridge, 2006). Beyond therapeutics’ insufficient brain exposure, the BBB also plays a major role in the underlying pathophysiology of many of these CNS disorders which are usually associated with vascular hyperpermeability, transporter deficiencies, or an increase in leukocyte adhesion molecules, resulting in an abnormal, uncontrolled movement of cells and neurotoxins across the BBB vessel walls (Pardridge, 2006).

For studies of barrier function and dysfunction, *in vivo* models are of highest physiological relevance since the BBB is embedded in its natural microenvironment. These models are, however, limited in their throughput. Furthermore, animal models may not predict BBB penetrance and efficacy of drugs in humans due to interspecies differences in the molecular composition of the BBB (Uchida et al., 2011). Deciphering the underlying molecular mechanisms and performing translatable real-time quantitative assessments of drug transport across the BBB, such as screenings for BBB-penetrant therapeutic antibodies, are therefore greatly limited in an *in vivo* setting. In contrast, *in vitro* BBB models offer faster, yet simplified approaches for targeted drug screening as well as for fundamental research, and importantly can be humanized to overcome translatability issues.

Human BBB organoids provide a model that enables maintaining endothelial cells in close juxtaposition. A limitation of this system, however, is that they essentially lack flow since microvessel-like structures cannot be formed in organoids, rather endothelium-lined spheres are generated which can negatively impact cellular viability (Urich et al., 2013). Traditional two-dimensional (2D) *in vitro* BBB models such as the Transwell system, in which endothelial cells are cultured on semi-permeable membranes, have extensively been used for cell-based high-throughput screening assays and for studying basic BBB characteristics such as barrier permeability and transepithelial/transendothelial electrical resistance (TEER) (Abbott et al., 1992; Biegel and Pachter, 1994; He et al., 2014). These simplified systems lack simulation of blood flow conditions and have proved to insufficiently recapitulate *in vivo* phenotypes including the expression of key junctional proteins (such as claudin-5) and transporters (such as Glut-1 and insulin receptor) (Campisi et al., 2018; Wevers et al., 2018). To overcome some of these limitations, several 3D microfluidic and organ-on-a-chip BBB models have been developed enabling co-culture and fluid flow (Prabhakarpandian et al., 2013; Herland et al., 2016; van Der Helm et al., 2016; Wevers et al., 2018; Oddo et al., 2019; Park et al., 2019). Nevertheless, a number of these models exhibit other limitations such as non-physiological rigid ECM substrates, failure to feature blood vessel-like geometry and the lack of controlled flow that resembles the hemodynamic forces which is known to be crucial for microvascular function (Herland et al., 2016). Hence, there is an essential need for *in vitro* BBB models that better mimic the brain microvessel environment including unidirectional flow, physiological shear stress, absence of artificial membranes, and presence of the cylindrical geometry typical of capillaries to facilitate the complex cell-cell interactions and physical ECM mechanics known to be intrinsic to the *in vivo* BBB. In order to be capable of providing molecular mechanistic insights, these models need to also be compatible with advanced imaging and live 3D tracking of labeled molecules.

Along these lines, we describe here our efforts to develop and use an *in vitro* human BBB-on-a-chip consisting of a 3D microfluidic model with a hollow brain-like microvessel in which a continuous monolayer of cells can grow at the interphase between the lumen and the underlying ECM. This system allows controlled unidirectional flow, within the lumen of the artificial microvessel, of media including substrates of interest, for instance drug candidate biologics. An important characteristic of our device is its open design that also allows direct access of reagents from the surrounding space to the underlying ECM. We demonstrate the utility of this open design of organ-on-a-chip model by showing it is amenable for quantitative 3D live fluorescence imaging using spinning disk confocal or lattice light sheet microscopy (LLSM) and for high resolution electron microscopy. This model is set out to provide insights into molecular mechanisms involved in the transcytosis of biologicals at extraordinary detail which will further support the development of antibody-shuttle technologies across the human BBB. Detailed imaging, for example, can be very useful to follow endo-lysosomal trafficking in real-time, informing on fate of antibodies and viruses when entering endothelial cells, thus informing on better designs of biologics and viral vectors that more efficiently penetrate or inhibitors for the transcytosis of pathological viruses.

## MATERIALS AND METHODS

### Cell culture

Human brain derived microvascular endothelial cells (TY10 cell line) were isolated from normal brain tissue from a patient with meningioma. Cells were immortalized with retroviral vectors harboring a SV40 large T antigen gene that is engineered to drive proliferation at 33 °C (Sano et al., 2010; Maeda et al., 2013; Sano et al., 2013) and have been used to model the BBB in previous studies (Takeshita et al., 2014; Karassek et al., 2015; Spampinato et al., 2015; Shimizu et al., 2017; Takeshita et al., 2017; Wevers et al., 2018). Cells were cultured at 33 °C, 5% CO_2_ in T75 flasks BioCoat (Corning, 354485, MA, USA). TY10 cells were used between passage 17–25 and cultured in ScienCell complete endothelial cell medium (ScienCell, 1001, CA, USA). Cell detachment was performed using Accutase® (Corning, 25-058-CI, MA, USA) when cells were ∼80–90% confluent before being seeded into the microfluidic devices. Cells were routinely tested for mycoplasma contamination and found negative.

### TY10 stably expressing eGFP

A lentiviral vector expressing a plasma membrane targeted eGFP (memGFP) containing a chimera of the N-terminal 41 amino acids of human myristoylated alanine-rich C-kinase substrate (MARCKS) fused to eGFP was made by co-transfection of a plasmid harboring memeGFP and Virapower Packaging Mix (Thermo Fisher, K497500, MA, USA) into 293T cells. Culture media was harvested 72 h later, cellular debris pelleted by low-speed centrifugation, and further clarified by 0.45 um filtration with Millipore steriflip vacuum filters (EMD, SLHV033RS). The supernatant from the viral preparation was added to a flask of TY10 cells during passaging (4 ml of viral supernatant preparation mixed with 4 ml of cell suspension were added to a T25 Corning BioCoat flask) and allowed to incubate for 24 h at 33 °C before switching back to the normal feeding schedule of every other day. The cells were sorted by flow cytometry for eGFP positive cells after 10 days in culture and subsequently expanded and maintained as described above.

### hmAb

The recombinant monoclonal human IgG1 antibody (produced by Biogen) was expressed in CHO cells and purified through Protein-A Affinity Chromatography. The purified protein was fluorescently labeled with Alex Fluor™ 568 and 647 protein-labeling kits (Thermo Fisher Scientific Cat# A10238 and A30009) to produce hmAb-AF568 and hmAb-AF647 following the manufacturer’s protocol.

### BBB-on-a-chip

#### Fabrication

Molds for microfluidic channels with a width, height and length of 5 mm, 0.16 mm and 13 mm, respectively, were designed with AutoCAD software (AutoDesk Corp., CA, USA) and produced by Outputcity (CAD/Art Services, Inc., Oregon, USA). Microfluidic devices were subsequently produced by soft lithography; Sylgard 184 elastomer Polydimethylsiloxane (PDMS) was mixed with curing agent (Sylgard 184 silicone elastomer kit, Dow Corning, Midlands, USA), at a 5:1 ratio in a mixer including a 2 min de-foaming step before pouring it onto the master silicon wafer designed by our lab and spin-coating at 400 rpm for 40 seconds. The utilized speed yielded a PDMS film of 160 µm thickness that was degassed in a vacuum desiccator for 10 min and cured in an oven at 65°C for 1h. The PDMS film was peeled off the master and placed in a plastic petri dish at 65°C overnight to fully cure the PDMS. Incomplete curing of PDMS leaves uncross-linked oligomers within the material that can leach out and contaminate the culture medium (Halldorsson et al., 2015). So prior to assembly, precut PDMS slabs containing the embossed chip microstructures and 5×5 mm end pieces were pre-cleaned by contact with Scotch tape and subjected to organic solvents to extract uncured PDMS-oligomers (Lee et al., 2003). PDMS slabs were incubated in a sealed jar on a rocker for 24h at room temperature (RT) in each of the following solvents and in this order: Triethylamine, Toluene, Ethyl Acetate, and Acetone (Sigma). Organic solvent was evaporated from the PDMS by incubation in a 100°C oven for 2h. Extracted PDMS remains hydrophilic for prolonged times after activation and prevents the leaching of uncured oligomers into the media (Lee et al., 2003; Kim and Herr, 2013). The PDMS extraction step has significantly improved the success rate of final chips in the downstream applications. The PDMS pieces of the chip were placed on a fluorinated ethylene propylene (FEP) sheet and exposed to air plasma at 700mTorr, 30 W for 1.5 min using Plasma Etcher (Harrick Plasma), bonded to the PDMS end pieces before punching 1mm holes for the tubing and plasma-bonding to the #1.5 glass coverslip that was pre-cleaned by incubation in isopropyl alcohol, acetone and 0.5M KOH for 30 min each in a sonication water bath, rinsing in dH_2_O water and blown dry with filtered nitrogen gas. We tested several glues to fix the Tygon microbore tubing, 0.010” x 0.030”OD (Cole-Parmer) on the chip and SLOW-CURE™ 30 min epoxy (Bob Smith Industries; BSI206, CA, USA) resulted in the sturdiest connection and further allowed us to remove air bubbles in the epoxy after mixing through centrifugation for 30 seconds at 14k g in a tabletop centrifuge. Following an overnight incubation at RT to fully cure the epoxy, the chip was activated by air plasma treatment as above and further cleaned by injecting sequentially 0.5 ml of each acetonitrile, purified water, 0.5M KOH and again dH_2_O. We functionalized the PDMS surface in a three-step process involving oxygen plasma treatment, amino-silane conjugation and glutaraldehyde derivatization. To functionalize both glass and PDMS surface with primary amine groups, the chips were silanized by immediately adding 0.5 ml of a fresh 1% aqueous solution of 3- (Ethoxydimethylsilyl) propylamine (Sigma, 588857) into the culturing chamber of the chip and incubated for 15 min at RT before rinsing with twice with 1 ml dH_2_O water. Subsequently, the surfaces were further functionalized by filling the devices with 2.5% glutaraldehyde (Electron Microscopy Services, 16200). After incubating for 15 min, the devices were rinsed extensively with deionized water. The Schiff bases formed on proteins after glutaraldehyde immobilization are stable without further reduction, as has been demonstrated in surface-protein conjugation (Kim and Herr, 2013).

#### Formation of lumen and collagen matrix

A Pluronic F-127 (Sigma, P2443) passivated 100 µm acupuncture needle was inserted from the outlet towards the inlet of the BBB-on-a-chip to provide the required scaffold for the culturing matrix as indicated in Fig 1C. The selected size of the acupuncture needle should prevent the leakage of unpolymerized collagen into the microfluidic channel of a smaller diameter (80 µm). A hydrogel consisting of extracellular matrix (ECM) proteins made of a final concentration of 7.0 mg/ml Type I rat tail collagen (Corning, 354249, MA, USA) was used in all experiments. To make 200 µL of hydrogel solution, 39 µl of Endothelial Cell Medium (ECM basal media with no FBS; 1001b, ScienCell, Carlsbad, CA, USA), 1 µl of a basic solution (1.0 N NaOH, Sigma) and 14 µl of 10X Ham’s F-12 (Thermo Fisher, 31765092, MA, USA) were added to 135 µl of the collagen I.

**Figure 1.**
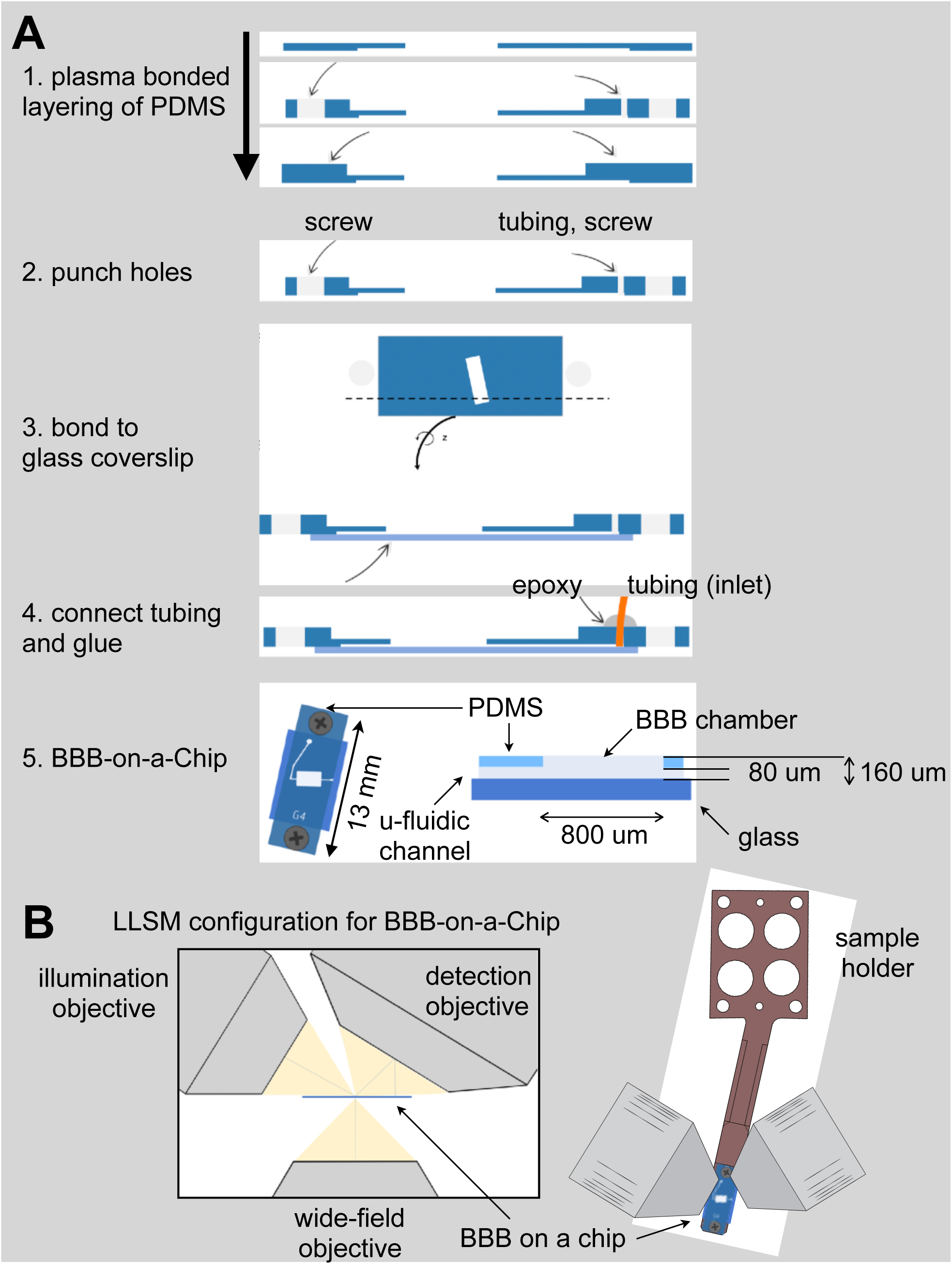
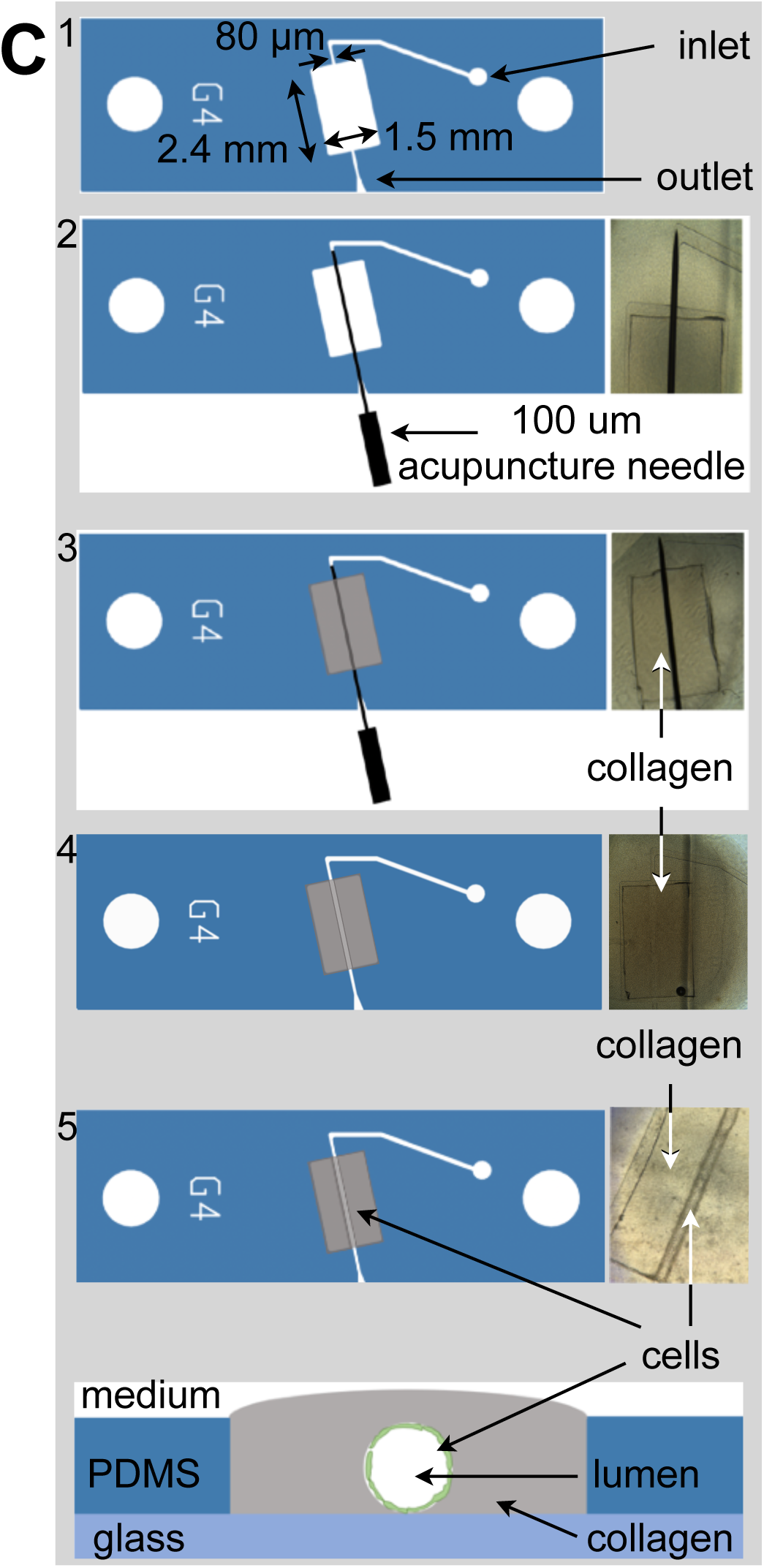
Fabrication of the BBB-on-a-chip. **(A)** Schematic representation of the sequential steps used to fabricate the BBB-on-a-chip. The open design permits direct access of aerated medium from the top to the collagen containing embedded cells while at the same time allowing flow of medium along the tubular channel with endothelial cells surrounding the boundary between the lumen of the artificial microvessel and the collagen. **(B)** Arrangement of the BBB-on-a-chip within the available optical path between the illumination and detection objectives of the lattice light sheet microscope. A sample holder is used to place the BBB-on-a-chip under the LLSM objectives and to facilitate fine adjustments of its position. **(C)** Steps used to create the artificial microvessel. An acupuncture needle of 100 µm diameter is placed between the inlet and outlet. After the collagen gelled, the needle was pulled, resulting in a void channel in which endothelial cells are seeded. The cross section on the bottom shows the final disposition of the endothelial cells (green) which line the inner surface of the collagen lumen, representing the artificial BBB microvessel.

We found that the collagen gel tended to delaminate from the PDMS culturing chamber as soon as we started the flow, and so we enhanced the standard collagen matrix protocol by adding Genipin^®^ (Sigma, G4796). Genipin is a crosslinking agent that covalently attaches to primary amino groups exposed on protein surfaces (Sung et al., 1998). Furthermore, Genipin monomers form covalent intermolecular crosslinks that in the case of a collagen matrix results in bridging adjacent fibers at points of contact (Yoo et al., 2011; Chan et al., 2014). Thus, to achieve a stiff and resilient collagen matrix we mixed collagen with Genipin prior pipetting it into the culturing chamber of the BBB-on-a-chip. The solution of 135 µl collagen, 39 µl ECM, 14 µl 10X Ham’s F-12, 1 µl 1N NaOH and 1 µl 20 mM Genipin was gently mixed and incubated on ice for a period of 5-10 min to get rid of any air bubbles which might generate during the mixing step, before being added to pre-chilled chips kept on ice for at least 15 min. The devices were subsequently incubated at 37°C to allow gel formation of the collagen matrix. Genipin improved the stability of the PDMS-collagen interaction such that delamination was never observed for up to 12 days. After removal of the acupuncture needle, non-reacted Genipin was quenched by covering the top of the collagen BBB-on-a-chip with PBS containing 1 mM Tris pH 8.0 in PBS in addition to flowing the same solution through the cylindrical lumen for 15 min at 1 µl/min. Chips were then washed with 3 ml of PBS alone (added to the top of collagen) and flow of PBS alone for 15 min at 1 µl/min. Prior to cell seeding (see below), a solution containing complete ECM medium was injected to the lumen for 15 min at 1 µl/min.

#### Cell seeding

To line the lumen with TY10 cells and generate a perfused microvessel-like structure, two strategies were used to ensure uniform cell seeding. In the first strategy (cell concentrator chip, Fig 2A), we designed a gravity-based microfluidic cell concentrator to reach a sufficiently high density of cells for seeding of the collagen lumen using minimal cell concentration. A PDMS chip whose single channel splits up into four microchannels that merge again into a single channel after 5mm was used as bottom layer with a central 2.5mm collection chamber. To securely fit a 25 mm long silicon tubing of OD 4mm / ID 2.5mm, a second PDMS layer with 4mm hole was bonded as a lid and the tubing was fixated with epoxy glue. The inlet of the concentrator chip was connected via tubing to a syringe pump and the outlet to the BBB-on-a-chip. TY10 cells were resuspended to 0.1 million cells/ml and transferred into a 1ml syringe. The cells settled by gravity within 15 min at the collection chamber on the bottom glass surface which was passivated with 0.01 mg/ml Poly-D-Lysine-PEG to prevent the cells from sticking to the glass. A plug compromised of an epoxy filled pipette tip was inserted into the central tubing to prevent upwards flow before initiating the flow. Applying flow through the microfluidic channel resulted in shear force that pushed the cell bolus into the tubing leading to the culturing chamber of the BBB-on-a-chip.

**Figure 2.**
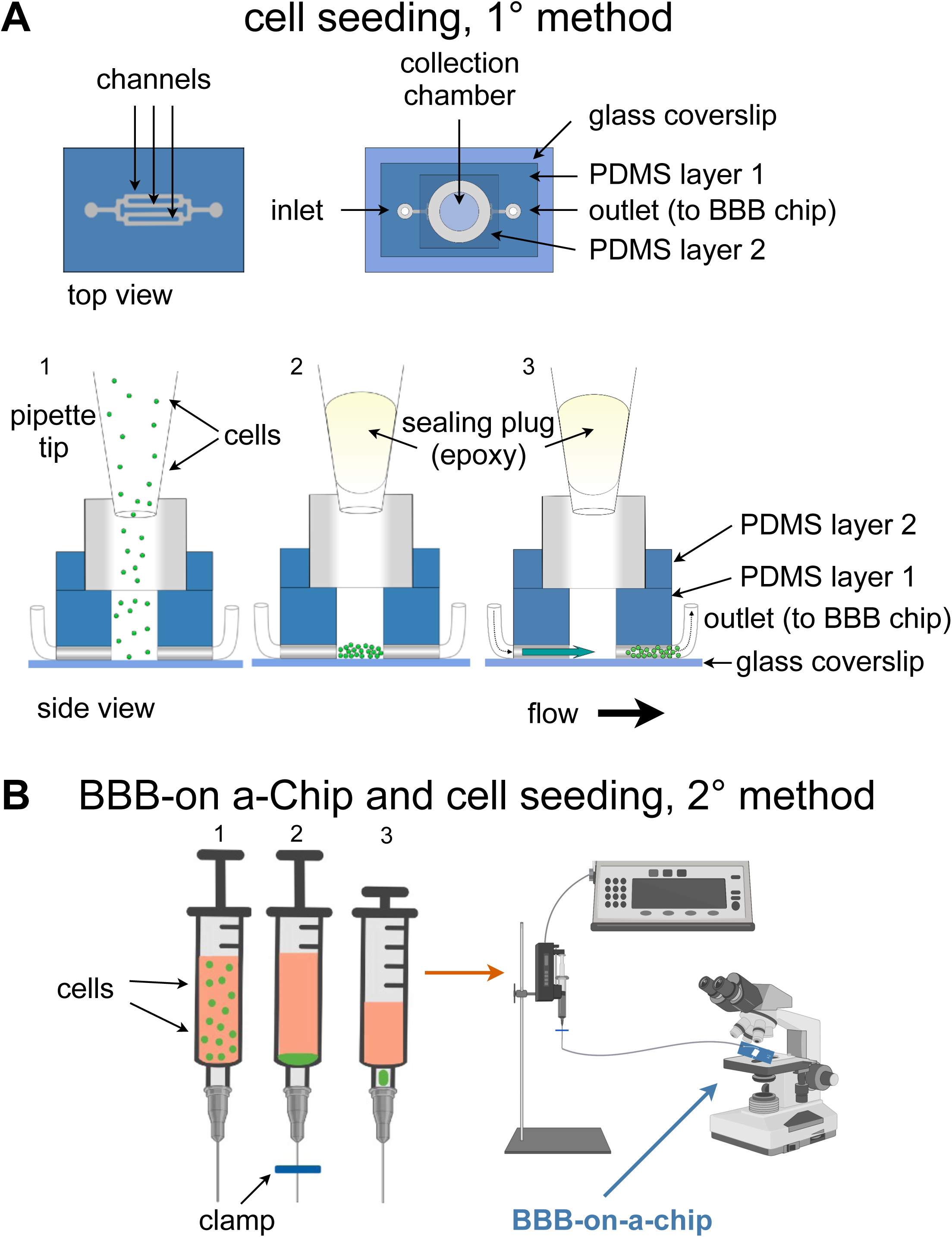
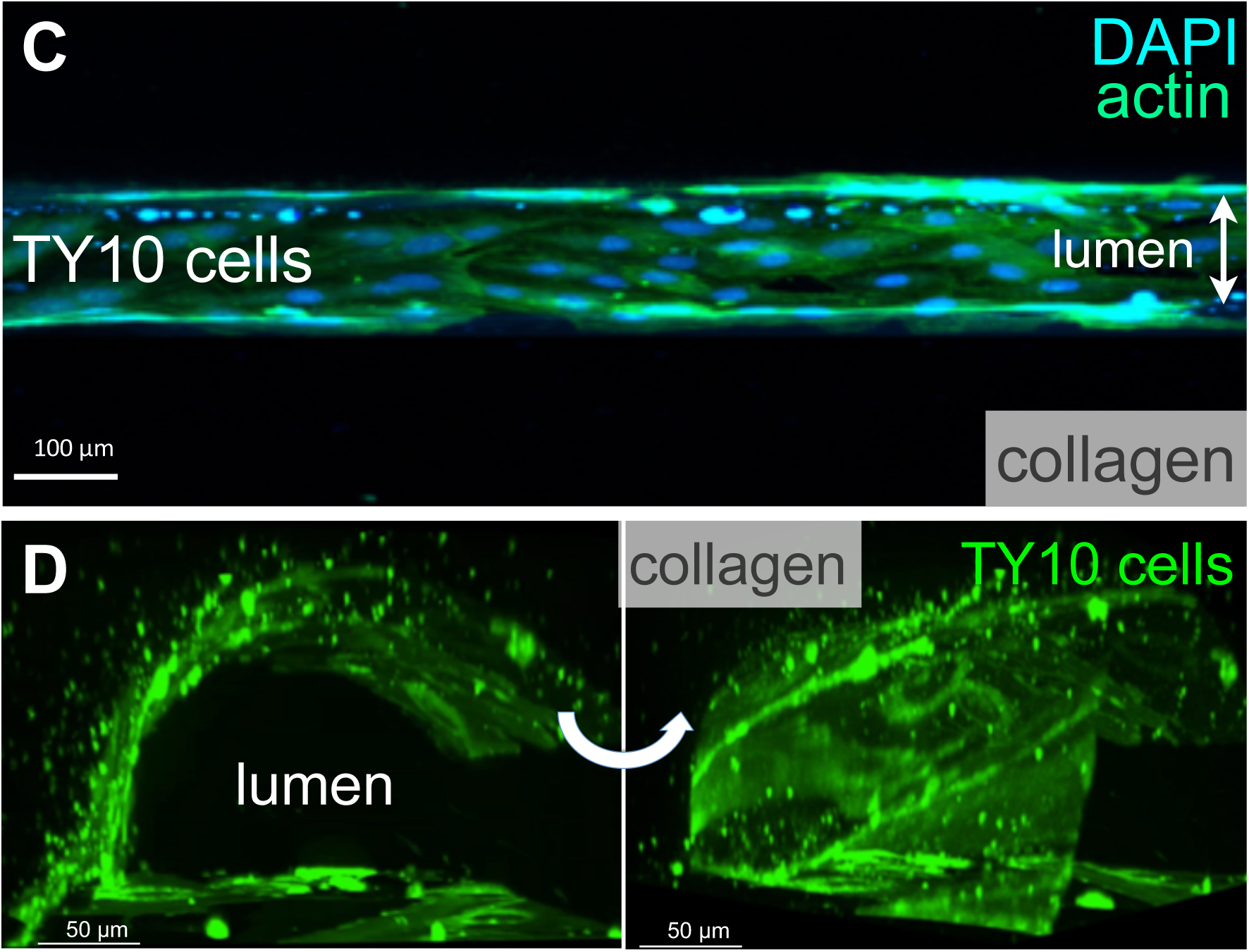
Cell seeding procedures and formation of a BBB microvessel using TY10 cells. **(A)** Cell seeding, 1^st^ method. Representation of the steps used to increase the cell concentration before injection into the BBB-on-a-chip. Cells are allowed to settle by gravity (left and central panels) in the cell seeder and then injected as a bolus into the BBB-on-a-chip. Prior to injection, the chamber is capped with a plugged pipette tip (central panel). **(B)** Cell seeding, 2^nd^ method. Representation of the steps used to inject cells into the BBB-on-a-chip in a more controlled and uniform manner compared to the 1^st^ method. Cells are allowed to settle at the bottom of a syringe, and are then delivered into the BBB-on-a-chip at constant flow controlled by a syringe pump. **(C)** Representative image of a chemically fixed sample of TY10 cells after they were grown in the BBB-on-a-chip with medium flowing at 1 µl / min for 7 days. Volumetric image was obtained using a spinning disk confocal microscope. Maximum z-projection is shown for a sample stained with DAPI (nuclei, blue) and an antibody specific for actin (green). Scale bar, 100 µm. **(D)** Representative volumetric image of a live sample of TY10-eGFP cells expressing membrane bound eGFP obtained using spinning disk confocal microscopy. The image highlights the organization of the cells as a monolayer at the boundary between the lumen of the artificial microvessel and the collagen scaffold. The cells on the bottom illustrate cells growing between the glass slide and the lumen. Panels are rotated 90-degrees from each other. Scale bar, 50 µm. See associated Movie 1.

In the second strategy a simplified procedure was implemented in order to enhance the experimental turnover as illustrated in Fig 2B and its results section. In brief, TY10 cells were harvested and resuspended to 1 million cells/ml. 900 µl of the cell solution was then mixed with 100 µl solution of collagen IV (Sigma, C5533), fibronectin (Sigma, F2518), and laminin (EMD Millipore, AG56P) at 5:1:1 concentration ratio before transferred into a 1 ml syringe with BD Luer-Lok (BD, 309628, NJ, USA). The syringe was then hanged vertically to allow cell settling by gravity for 10-15 min with no flow. Cell seeding was initiated under flow for about 15-20 min at 1 µl/min. Cell seeding was monitored by visual inspection using a microscope to observe the BBB-on-a-chip placed inside a petri dish kept under sterile conditions.

After either procedure, chips were incubated at 37°C for a minimum of 4h before being perfused with fresh ECM media via positive pumping to wash out the unattached cells. The chips were maintained under continuous unidirectional flow at a rate of 1 µl/min in a cell culture incubator at 37 °C, 5% CO_2_. Confluent TY10 monolayers were formed typically after 72h and the BBB-on-a-chip devices were used for subsequent analyses following 7 days after seeding incubation in all experiments.

### Immunostaining

The collagen samples including the lumen were fixed and permeabilized using BD Cytofix/Cytoperm Kit (BD, 554714, NJ, USA) according to the manufacturer’s protocol. In brief, Cytofix was added to a 1 ml syringe and perfused through the microvessel at a flow rate of 1 µl/min for 30 min before changing to the Cytoperm buffer (diluted 1:10 in PBS) for 1h at 1 µl/min. The flow was then stopped and the BBB-on-a-chip placed on a well of a six well plate in 2 ml of Cytoperm buffer (diluted 1:10 in PBS) overnight. The microvessel was stained with AlexaFluor488 Phalloidin (Thermo, A12379, at 1:100 dilution) and NucBlue (Thermo, R37606, 1 drop/500 µl) by perfusing the dyes in Cytoperm buffer at 1 µl/min through the microvessel for 1h, followed by washing with 0.1% BSA in PBS buffer for 4h at 1 µl/min. Images were collected using a 3i spinning disk microscope (40x water immersion objective) through the bottom glass slide.

### Barrier integrity assay

Chips were washed with ECM culture medium (once with 1 ml added to the top of collagen followed by a flow at 1 µl/min for 15 min) to ensure proper flow profiles during the subsequent barrier integrity assay. Next, all medium was aspirated from the chip and 1 ml of medium without fluorescent compound using Gibco® FluoroBrite™ DMEM (Thermo Fisher, A1896701, MA, USA) was added. Medium containing 25 µg/ml of 10 kDa FITC-dextran (Sigma, FD10S) or hmAb-AF568 was added through the inlet at a flow rate of 1 µl/min for all the experiments. The inlet was connected to microvessels with and without TY10 cells and image acquisition was started. Leakage of the fluorescent substrate (dextran or hmAb) from the lumen of the microvessel into the adjacent collagen matrix was imaged using a spinning disk confocal microscope with 40x water immersion objective. The fluorescence intensity profiles and ratios between the fluorescent signal in the basal and apical region of the microvessel tube were analyzed using MATLAB (MathWorks, MA, USA). Apparent permeability (Papp) was used for quantifying diffusional permeability as described (Yuan et al., 2009). In brief, Papp was calculated by analyzing total fluorescence intensity in the imaged 2D area of the lumen and collagen and then applying Papp = (1/ΔI) (dI/dt)_0_ (r/2), where ΔI is the increase in total fluorescence intensity upon adding labeled dextran or labeled hmAb to the lumen, (dI/dt)_0_ is the temporal initial rate of linear increase in intensity as the labeled molecules diffuse out of the microvessel into the surrounding collagen matrix, and (r) is the radius of the microvessel (100 µm for our BBB-on-a-chip). All experiments were carried out at n = 4–6; exact numbers are mentioned per experiment in figure captions. Graphs were plotted using GraphPad Prism 6 (GraphPad Software, San Diego, CA, USA).

### Spinning disk confocal imaging

Imaging was done using a Marianas spinning disk confocal microscope (3i, Denver, Colorado) with the water immersion objective lens LD C-Apochromat 40x/1.1 (Carl Zeiss, Jena, Germany). The images consisted of 512 × 512 pixels with a pixel size of ∼ 333 nm. The EMCCD camera settings (gain, speed, intensification, and exposure) and laser power were maintained throughout the imaging experiments. Images were acquired using SlideBook 6 (3i, Denver, Colorado) and data analysis carried out using SlideBook 8 and custom made software using MATLAB 2017A (Natick, Massachusetts). For the analysis of heat maps and Papp, ROIs were the entire original field of view.

### LLSM imaging

A BBB-on-a-chip fabricated on 8*5-mm rectangular #1.5 glass coverslip was picked with forceps, and placed in the sample bath of the 3D LLSM. The sample was imaged in a time series in 3D using a dithered multi-Bessel lattice light-sheet by stepping the sample stage at 200 nm intervals in the s-axis equivalent to ∼104 nm translation in the z-axis. Each 3D stack corresponded to a pre-deskewed volume of ∼80 µm × 120 µm × 47 µm (800 × 1200 × 451 pixels). The sample was excited with a 488-nm laser (∼100mW operating power with an illumination of ∼77 µW at the back aperture), a 560-nm laser (∼100mW operating power with an illumination of ∼176 µW at the back aperture) and a 642-nm laser (∼100mW operating power with an illumination of ∼121 µW at the back aperture) to acquire 451 imaging planes, each exposed for ∼44.1 ms and recorded with two Hamamatsu ORCA-flash 4.0-V2 cameras; thus, each 3D image took ∼60 s to acquire. The inner and outer numerical apertures (NAs) of excitation were 0.513 and 0.55, respectively. The overall 3D volume of ∼240um × 880um × 180um was obtained by stitching together 165 (3 × 11 × 5) 3D stacks, with an overlap of 40 µm and 9 µm in the y-axis and z-axis respectively.

### Transmission Electron Microscopy

The collagen matrix including the lumen was washed 3 times with PBS and then fixed by immersion in 5 ml of fixing solution (2.5% glutaraldehyde, 2% sucrose, 50 mM KCl, 2.5 mM MgCl_2_, 2.5 mM CaCl_2_ in 50 mM Cacodylate buffer pH 7.4) (Sigma) and kept at 4°C, overnight in the dark. Fixed collagen samples were washed 3 times with a solution containing 50 mM PIPES pH 7.4 (Sigma, P6757) kept in ice and then they incubated for 2h in ice and in the dark in a freshly prepared staining solution (SSI) made of 1% OsO4 (Electron Microscopy Sciences, 19190), 1.25% potassium hexacyanoferrate (II) (Sigma, 455989) and 100 mM PIPES pH 7.4. Samples were rinsed 3 times with ice-cold water and incubated again for a second time for 30 min in ice and in the dark with a freshly prepared staining solution II (SSII). SSII was prepared by 1:100 dilution of SSI in a freshly prepared 1% thiocarbohydrizide (Electron Microscopy Sciences, 21900). Finally, the samples were washed for 3 times with ice-cold H_2_O and then incubated overnight in the dark in 1% uranyl acetate (Electron Microscopy Sciences, 22400) at 4°C.

For the dehydration and embedding step, the fixed and stained samples were first washed 3 times with ice cold water and then subjected to dehydration with a 20-50-70-90-100% ethanol – 100% acetone dehydration series. Samples were then infiltrated overnight, at 4°C with 50-50 acetone-Epon812 epoxy resin (Electron Microscopy Sciences, 14120). Next day, the samples were washed 3 times with 100% Epon812 and then kept in an oven for 36 h at 60°C. Sections of 60-70 nm were cut transversally to the long axis of the lumen and imaged with a JEOL JEM 1200 EX TEM microscope with a voltage of 80 KV and a nominal magnification of 15,000.

### Statistical analysis

StatsDirect 3 (Liverpool, UK) was used for one-way ANOVA and student’s t-test analyses with Bonferroni *post-hoc* correction. Data were presented as mean±standard error of the mean (S.E.M.) where (***) denotes a statistically significant difference with p< 0.0001 and (ns) indicates a statistically non-significant difference.

## RESULTS AND DISCUSSION

We designed an open design microfluidic chip to generate a 3D microphysiological model of the human BBB readily accessible to optical imaging. The unique characteristics of this novel BBB-on-a-chip are the open design of the cell culture chamber and a cylindrical hollow lumen amenable to continuous flow within a casted gel of extracellular matrix (ECM) components. The open system allows direct access of the collagen matrix for efficient exchange of gases and medium in addition to readily access to optical imaging while cells growing with continuous unidirectional flow at the interphase between the casted gel and the lumen mimic the environment of a microvessel.

Fig. 1A graphically summarizes the sequential steps used to build our BBB-on-a-chip model. It is based on sequential bonding using soft lithography (Bischel et al., 2013) of thin layers of optically clear PDMS on top on a glass microscope slide. The geometry and dimensions of the BBB-on-a-chip were optimized for its use with three major complementary forms of live 3D optical imaging, spinning disk confocal, lattice light sheet microscopy (LLSM) and the recently developed variant, LLSM modified with adaptive optics (AO-LLSM) (Gao et al., 2019). We chose to include access to LLSM and AO-LLSM because these imaging modes have revolutionized fluorescence optical microscopy providing volumetric imaging with unprecedented high spatial and temporal precision with minimal bleaching and phototoxicity (Liu et al., 2018). Spinning disk confocal microscopy is performed through the glass slide at the bottom of the BBB-on-a-chip, while LLSM or AO-LLSM are carried out from the open top as illustrated in Fig. 1B. The device is also suited for chemical fixation and the sample preparation required for high-resolution electron microscopy visualization.

The consecutive stages used to generate the cylindrical hollow lumen within the casted gel followed by seeding of endothelial cells on the wall of the lumen are depicted in Fig. 1C and described in detail in methods. It involved first placing an acupuncture needle between the microfluidic inlet and outlets (Fig. 1C, 2) followed by casting a collagen matrix (Fig. 1C, 3), gentle removal of the needle after collagen gelation (Fig. 1C, 4) and ending with cell seeding under flow (Fig. 1C, 4). A Pluronic F-127 passivated 100 µm acupuncture needle was inserted from the outlet towards the inlet to provide the required scaffold for the culturing matrix. Collagen type I (7 mg/ml) has been used to assemble the culturing scaffold. The chosen diameter of the needle enables the device to recreate artificial microvessels where an endothelial monolayer is formed against a collagen matrix and is stably maintained by surface tension and shear stress.

Cell seeding was done with two methods. The first one involved use of a cell-concentrator chip designed and operated as indicated in Fig 2A. Cells in suspension were placed on a pipette tip linked to the top of the cell-concentrator, allowed to settle by gravity for 15 min to a cell density of ∼0.1 million/ml, and cells then injected into the BBB-on-a-chip with the aid of a syringe pump. Before activation of the syringe pump, we replaced the pipette feeding cells with an epoxy-plugged pipette as a way to prevent backflow. Afterwards the bolus with cells reached the BBB-on-a-chip, flow was then stopped allowing cells to settle for 24 hrs so they could attach to the internal walls of the cylindrical lumen of the BBB-on-a-chip. They were grown for 7 days under flow, at which point they established a monolayer and hence were ready for the imaging experiments. The second, and preferred cell seeding method (Fig. 2B), involved use of a 1 ml syringe driven by a mechanical syringe pump. A solution containing ∼ 900 µl of 1 million/ml cells in medium mixed with 100 µl of a solution containing laminin 1, fibronectin and collagen type IV, was placed in a vertically oriented syringe and cells allowed to settle for ∼10-15 min. Afterwards, at flow of 1 µl/min was applied for 10-15 min in order to inject the cells into the BBB-on-a-chip; cells were then allowed to settle within the lumen of the BBB-on-a-chip and attach to the internal walls of the lumen for 4 hrs at 37°C. As with the first method, cells were then grown for 7 days at a flow rate of 1 µl/min, before their use for imaging. This simpler cell seeding method is particularly advantageous for cases in which the cellular supply might be limited such as when using primary cells or iPSC-derived cells from patients. Extent of seeding was optically monitored with the aid of an inverted microscope by direct inspection of the BBB-on-a-chip placed inside a closed petri dish to ensure sterility.

In the brain, the basement membrane in contact with the endothelial cells of the BBB is comprised of fibronectin, laminin (Aumailley et al., 2005) and collagen type IV (Hartmann et al., 2007). Indeed, *in vitro* monolayers of endothelial cells grown on a matrix containing fibronectin, laminin 1, and collagen type IV exhibit enhanced TEER, suggesting a role for these molecules in promoting the formation of tight junctions (Tilling et al., 1998; Tilling et al., 2002; Gautam et al., 2016). To mimic the physiological BBB microenvironment and presumably also to enhance the seeding efficiency in the BBB-on-a-chip, we injected before cell seeding a solution containing fibronectin and laminin 1 for 30 min at 1 µl/min. TY10 cells were allowed to settle for 24 hrs at 37°C before starting the flow at 1 µl/min.

One of the major challenges of developing physiologically relevant *in vitro* BBB models is the availability of suitable brain-derived cells of endothelial origin and of human origin in particular. Primary human brain endothelial cells or cells differentiated from induced pluripotent stem cells (iPSC) derived from control or diseased patients are preferred for *in vitro* BBB models. Use of primary cells is restricted to very low passage numbers to prevent down-regulation of the unique features of the BBB (Reichel et al., 2003). More general, the difficulties in collecting and purifying these cells can considerably limit their use and reliability, as well as reproducibility ADD a ref specific for iPSCs (Bernas et al., 2010). Immortalized brain-derived cell lines can have great advantages such as accessibility and convenience of use especially for optimization purposes despite that some of the available lines might not exhibit all BBB characteristics (Kuhnline Sloan et al., 2012; Eigenmann et al., 2013; Wong et al., 2013). Nonetheless, certain cell lines may still exhibit the required properties for some pathophysiological and medicinal applications in a fit-for-purpose approach. We therefore chose immortalized human brain derived TY10 capillary endothelial cells that have been used to model the human BBB in a number of previous studies (Takeshita et al., 2014; Spampinato et al., 2015; Idris et al., 2018; Wevers et al., 2018).

TY10 cells are immortalized and proliferate at 33C, and stop growing and acquire a phenotype of primary brain endothelial cells at 37C (Maeda et al., 2013). These cells show a spindle-shaped morphology (Sano et al., 2013) and have well characterized barrier-forming features of endothelial cells, expressing the majority of essential tight junctional proteins, such as claudin-5, occludin, zonula occludens (ZO)-1 and ZO-2, as well as expression of P-glycoprotein irrespective of passage number (Sano et al., 2010). This is an important characteristic since shear stress is known to play a role in regulating signaling cascades (Conway and Schwartz, 2012), enhancing the expression of key genes associated with transporters and junctional proteins (Cucullo et al., 2011), and plays a pivotal role in BBB regulation (Neuwelt et al., 2008; Neuwelt et al., 2011).

As depicted in the representative fluorescence microscopy image of a chemically fixed sample stain for actin and DNA shown in Fig. 2C, TY10 cells grew as a monolayer at the interphase between the cylindrical lumen and the collagen matrix in the BBB-on-a-chip; grown for 7 days with constant flow at 1 µl/min of media, they appeared elongated along the flow axis, in agreement with previous findings (Ohashi and Sato, 2005; Aird, 2007). Further confirmation for the cell organization was obtained using TY10 cells stably expressing soluble eGFP grown in a similar way and then imaged by live cell fluorescence microscopy 3D imaging (Fig. 2D and related movie 1).

We further characterized, at the ultrastructural level, the TY10 cells grown under flow in our BBB-on-a-chip as a way to detect presence of tight junctions between adjacent TY10 cells grown at the lumen-collagen interphase by using transmission electron microscopy (TEM) (Fig. 3). The representative images highlight presence of a narrow gap between adjacent cells and occurrence of tight junctions. Similar images were observed along most cell-cell contacts between TY10 cells imaged visualized in other regions from this and other TEM sections.

**Figure 3.**
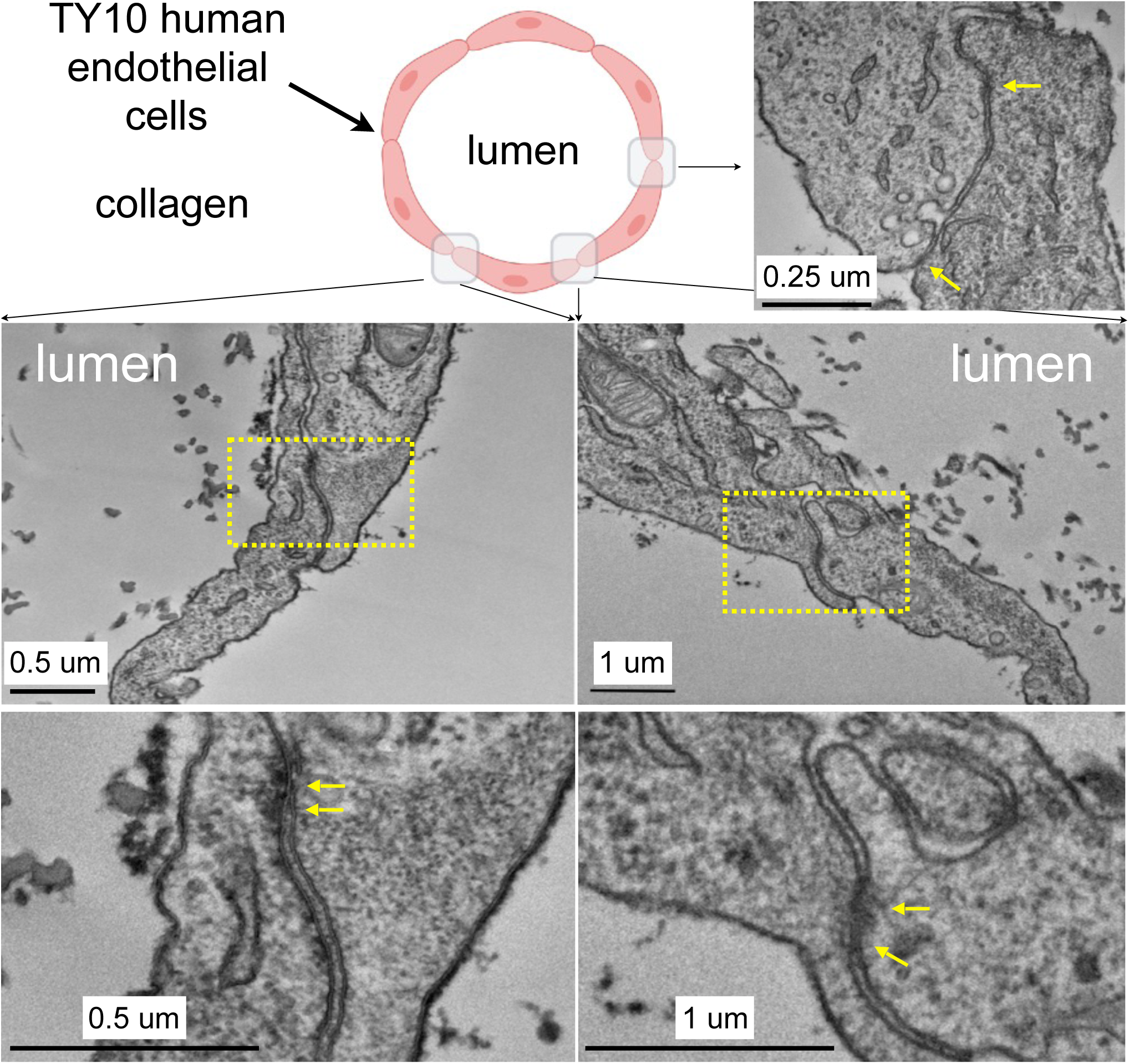
Transmission electron microscopy highlighting the appearance of junctions between TY10 endothelial cells within the BBB-on-a-chip. The upper left panel shows a schematic cross section of the BBB-on-a-chip highlighting the location of the monolayer of TY10 endothelial cells at the interface between the lumen of the microvessel and the collagen. The representative images in the bottom panels derive from junctions between opposite ends of adjacent cells. The yellow arrows highlight electron-density characteristic of tight junctions. Scale bars with corresponding magnifications are indicated.

TEER is frequently used to evaluate the integrity of the tight junctions and barrier function of *in vitro* models of BBB. Use of this approach is not practical for our BBB-on-a-chip model because the geometric constrains prevents us from positioning electrodes on opposite sides of the endothelial monolayer between the cylindrical lumen and the collagen matrix. To circumvent this limitation, we capitalized on our ability to use fluorescence optical imaging with our device as a way to determine permeability coefficient across the endothelial monolayer and hence establish the extent of the barrier function of cells grown in the BBB-on-a-chip. Using spinning disk confocal fluorescence microscopy, we determine the rate of transport of fluorescently tagged humanized monoclonal hmAb-AF568 (Fig. 4A, C, D) or 10 kDa FITC-dextran (Fig. 4B) across the lumen-matrix interphase in the absence or presence of cells. Unexpectedly, in the absence of cells, we found retention of hmAb-AF568 at the lumen-matrix interface (Fig. 4A, central fluorescence image). This retention appeared to be due to capture of the antibody by unreacted Genipin (the stabilizing primary amine crosslinker used to stabilize the collagen matrix, see Materials and Methods). Quenching the unreacted reagent (see materials and methods) fully prevented the hmAb-AF568 capture (Fig. 4A, right fluorescence image).

**Figure 4.**
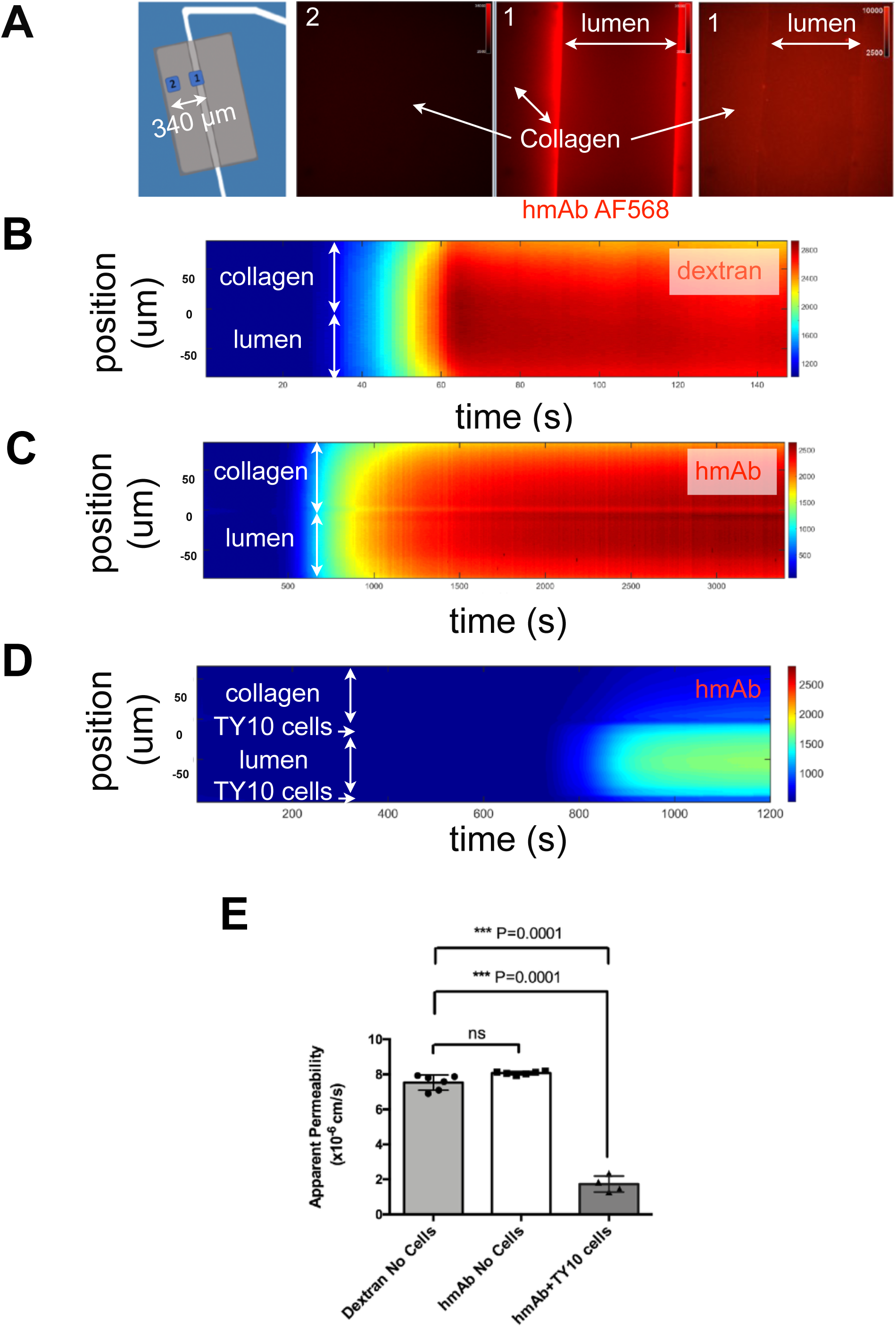
TY10 cells establish a functional barrier in the BBB-on-a-chip. **(A)** The left panel is a schematic representation of the experimental setup used to determine the apparent permeability of fluorescent solutes diffusing between the lumen of the artificial microvessel and the collagen in the absence and presence of TY10 cells. The boxed numbered areas represent typical regions imaged using spinning disk confocal microscopy. The fluorescent images are of hmAb-AF568 (red) applied at constant flow (1 µl /min) for 20 min in the absence (left and central panels) and presence (right panel) of 1 mM TRIS in addition to Genipin (a chemical crosslinker) added to stabilize the collagen matrix (see Methods). Genipin followed by 1 mM TRIS treatment dramatically decreased the non-specific retention of the antibody by the collagen. (**B-D**) Heat map representation of the fluorescence intensity of 10 kDa FITC-Dextran (**B**) or antibody hmAb-AF568 (**C**) from the lumen through the collagen as a function of time obtained at a flow of 1 µl /min. The significant decrease in the amount of antibody that passes through the endothelial cell layer is highlighted in panel (**D**), demonstrating that TY10 cells form a functional barrier in the BBB-on-a-chip. See associated Movies 2, 3 and 4. (**E**) Apparent permeability of data obtained from experiments carried as described in **(B-D)** for 10 kDa FITC-Dextran or hmAb-AF588 diffusion across the boundary between the lumen and collagen without cells (n=6 each), and for hmAb-AF588 diffusion with TY10 cells (n=4). Kruskal-Wallis with Conover-Inman post hoc tests were used to identify significant differences between samples. Error bars indicate S.E. M.; *** p≤ 0.001, ns: non-significant.

As shown by the time-dependent heat maps depicted in Figs. 4B and 4C, hmAb-AF568 and 10 kDa FITC-dextran freely diffused from the lumen towards the collagen matrix. Their apparent permeability, determined as the flux through a given unit area under gradient concentration (cm s^−1^) were also similar (Fig. 4E) as expected for molecules with comparable radius of gyration (1.86 nm and 5–6 nm, respectively) (Armstrong et al., 2004; Hawe et al., 2011). In contrast, presence of the TY10 monolayer at the lumen-matrix interphase significantly hindered the transport of hmAb-AF568 (Fig. 4D), with a significantly lower apparent permeability (Fig. 4E). We conclude from these observations that theTY10 cells grew as a relatively tight diffusion barrier in our BBB-on-a-chip, in agreement with similar previous results obtained with these and other cells using different *in vitro* human brain endothelial cell models (Eigenmann et al., 2013; Wevers et al., 2018).

As a final proof-of-principle, we verified the large-scale 3D optical fluorescence imaging capability of the BBB-on-a-chip, by using our LLSM set up (Fig. 5A) to visualize diffusing fluorescently-tagged antibodies captured by beads that had been embedded in the matrix; the antibody reached the beads upon diffusion from the lumen to the collagen. The result from one such experiment, using 3 µm SPHERO™ goat anti-human IgG coated polystyrene beads and hmAb-AF647 labeled antibodies is illustrated in Fig. 5B (arrows).

**Figure 5.**
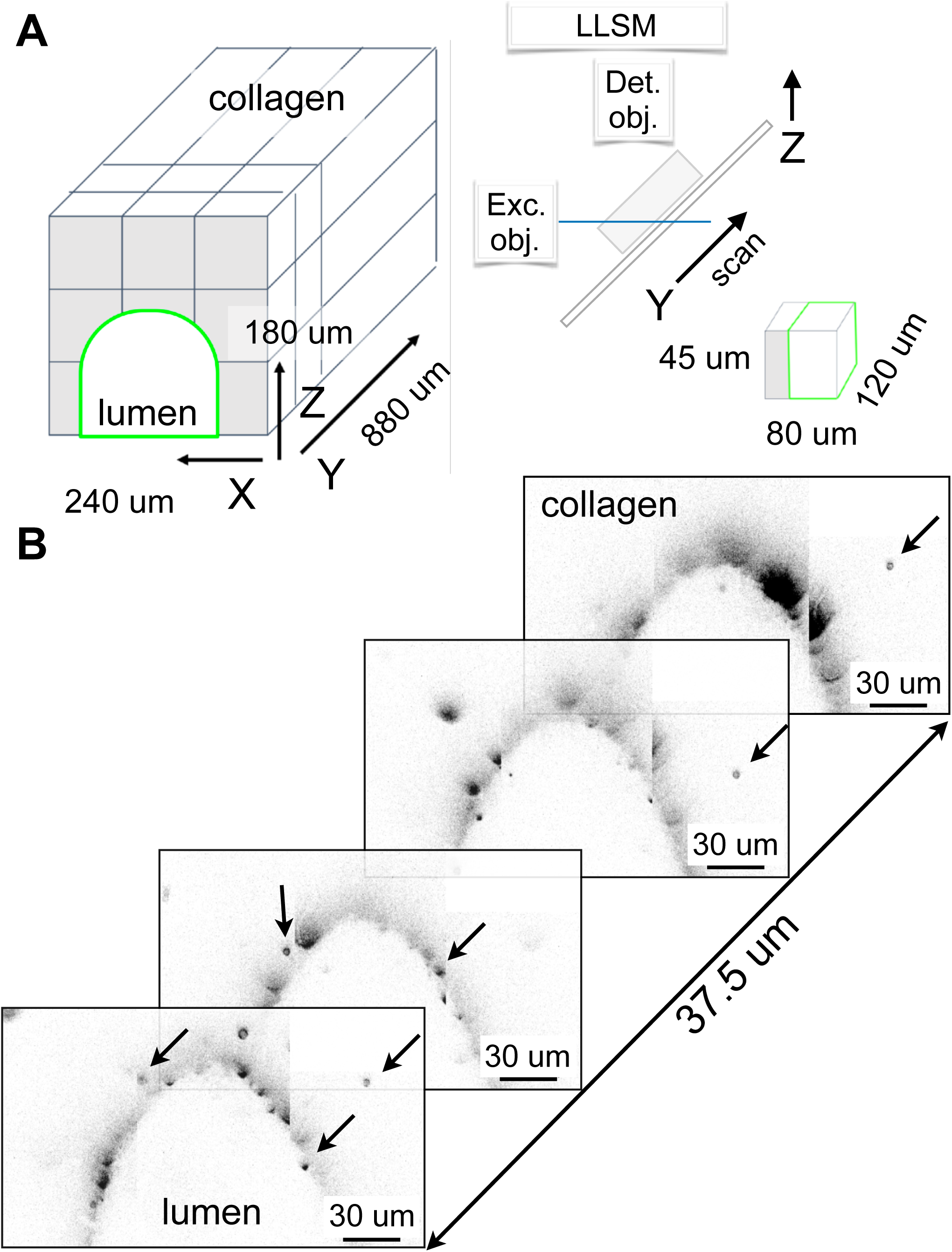
Visualizing IgG antibody diffusion in the BBB-on-a-chip device using LLSM. **(A)** Schematic representation of the volumetric imaging strategy used to visualize the lumen of the microvessel and surrounding volume within the BBB-on-a-chip using LLSM. The cubes represent the adjacent regions imaged by serial scanning of the sample using a distance of 100 nm between planes. **(B)** Selected planes corresponding to the volumetric imaging obtained using LLSM of a sample containing 3 µm SPHERO™ Goat anti-Human IgG coated polystyrene beads embedded in the collagen matrix. Before imaging, a solution containing hmAb-AF647 was perfused at 1 µl / min for 20 min. The fluorescent spots marked by arrows in the selected planes located 7.7, 13.7 and 16.2 µm apart correspond to the signal of hmAb-AF647 captured by the beads within the collagen matrix. Scale bar, 30 µm.

These results provide a supporting evidence showing how the intrinsic open design of our microfluidic BBB-on-a-chip device can be used to facilitate future studies of BBB physiology at a subcellular level, particularly since cells can be grown under controlled unidirectional flow conditions. The readily imaging accessibility of our BBB-on-a-chip is particularly suited for investigations of molecular transport mechanisms involved in the transcellular transport of biologicals, viruses or nanoparticles with extraordinary level of detail. Although we exemplified here its implementation with a LLSM system, its design also allows its use with AO-LLSM, which enables capture of high-resolution 3D movies of collective behavior of cells in a multicellular environment (Ji, 2017; Gao et al., 2019). Moreover, the open design of our BBB-on-a-chip facilitates use of chemical fixation of the biological material located within the collagen matrix and permits ease use of embedding procedures such as manipulations associated with conventional transmission microscopy, with high resolution volumetric imaging using Focused Ion Beam Scanning Electron Microscopy (FIB-SEM) and also with the newly developed modality of expansion microscopy combined with LLSM (Ex-LLSM) (Xu et al., 2017; Gao et al., 2019; Wassie et al., 2019).

## ACKNOWLEDGMENTS

We thank Fang Qian and Benjamin Smith (Biogen) for generating the fluorescent hmAb monoclonal antibody, G. Campbell Kaynor (Biogen) for providing the lentiviral construct expressing memEGFP, Ramiro Massol and Chris Ehrenfels (Cellular Imaging Unit, Biogen), Tegy John Vadakkan (T.K. Lab) for assistance with fluorescence microscopy imaging and Robin Kleiman (Biogen) for critical feedback.

## AUTHOR CONTRIBUTIONS

M.S., M.D., G.M. and T.K. conceived the project. M.S. and G.M. were involved in all experiments. M.D., M.S., G.M., S.U. and T.K. designed the BBB-on-a-chip. R.H. built BBB-on-a-chip devices under I.K. and M.S. supervision. G.M. and B.O. provided advice on all aspects of the cell biology associated with this project. I.K. and M.S. performed spinning disk confocal microscopy. K.G.H. performed lattice light sheet microscopy under T.K.’s supervision. G.D.C. helped with data analysis. G. d. N. performed electron microscopy. F.S., Y.S. and T.K. provided the parental TY10 cells and advised on cell culture procedures. T.K. supervised the project. M.S, B.O. and T.K. wrote the manuscript with comments from all authors.

## DATA AVAILABILITY

The datasets used and/or analyzed during the current study are available from the corresponding author T.K. upon request.

---

**Movie 1. Live cell imaging of a BBB-on-a-chip.** 3D rendition of a live sample of TY10-eGFP cells expressing membrane bound eGFP grown as a monolayer at the boundary between the lumen of the artificial microvessel and the collagen scaffold within the BBB-on-a-chip. The volumetric image was obtained using spinning disk confocal microscopy.

**Movie 2. Absence of a functional barrier between the lumen and the collagen matrix of the BBB-on-a-chip.** Time series of the fluorescence intensity presented as a heat map of 10 kDa FITC-Dextran diffusing from the lumen through the collagen as a function of time obtained at a flow of 1 µl /min. Data was obtained in the absence of a cell monolayer at the boundary between the lumen of the artificial microvessel and the collagen scaffold within the BBB-on-a-chip.

**Movie 3 TY10 cells establish a functional barrier in the BBB-on-a-chip.** Time series of the fluorescence intensity presented as a heat map of antibody hmAb-AF568 diffusing from the lumen through the collagen as a function of time obtained at a flow of 1 µl /min. Data was obtained in the presence of a monolayer of TY10 cells at the boundary between the lumen of the artificial microvessel and the collagen scaffold within the BBB-on-a-chip.

**Movie 4. TY10 cells establish a functional barrier in the BBB-on-a-chip.** Time series of the fluorescence intensity presented as a heat map of antibody hmAb-AF568 diffusing from the lumen through the collagen as a function of time obtained at a flow of 1 µl /min. Data is a second example obtained in the presence of a monolayer of TY10 cells at the boundary between the lumen of the artificial microvessel and the collagen scaffold within the BBB-on-a-chip.

